# Long non-coding RNA Cerox1 targets components of the mitochondrial electron transport chain to regulate the memory impairment caused by sleep deprivation

**DOI:** 10.1101/2025.09.15.676326

**Authors:** Karthick Ravichandran, Pallabi Kisku, Lakhuhong Ningchangmi, Premkumar Palanisamy, Stefan Strack, Ted Abel, Sourav Banerjee

## Abstract

Sleep deprivation (SD) impairs long-term memory, but the molecular mechanisms underlying the impact of sleep loss on memory are poorly understood. Molecular changes driven by SD have thus far focused on transcription and translation. Long non-coding RNAs (lncRNAs), a class of regulatory RNAs, have recently been recognized as an important player in memory research. However, it remains unclear how sleep deprivation modulates the expression of lncRNAs or their targets to lead to memory impairment. In this study, we explored the role of lncRNAs in the disruption of spatial memory caused by SD. We examined a set of synapse-associated lncRNAs that were identified through a transcriptome analysis after SD. Among them, we discovered that the lncRNA Cerox1 is downregulated in dorsal hippocampus following SD, and its levels recover after 2.5 hours of rebound sleep. Sleep is critical for the regulation of metabolism and sleep loss impairs mitochondrial function. Both sleep deprivation and Cerox1 knockdown were found to reduce complex I activity of the mitochondrial electron transport chain. This reduction of complex I activity is linked to the decrease in expression of a subset of complex I subunits including Ndufs1, Ndufs3, Ndufa3 and Ndufs6. Overexpression of Cerox1 has the opposite effect, leading to increased complex I activity. Sleep deprivation reduced ATP levels in the dorsal hippocampus, while Cerox1 overexpression restored it. SD disrupted memory consolidation, and this impairment was rescued when Cerox1 was overexpressed. Cerox1 transcript contains multiple miRNA binding sites that regulate the activity of the lncRNA. Notably, overexpression of Cerox1 transcript lacking miRNA binding sites did not rescue the memory deficit caused by SD. Our findings demonstrated that the impairment of memory consolidation after SD is linked to lncRNA-mediated control of mitochondrial electron transport chain activity essential for sustaining energy requirements.

**Graphical Abstract:** 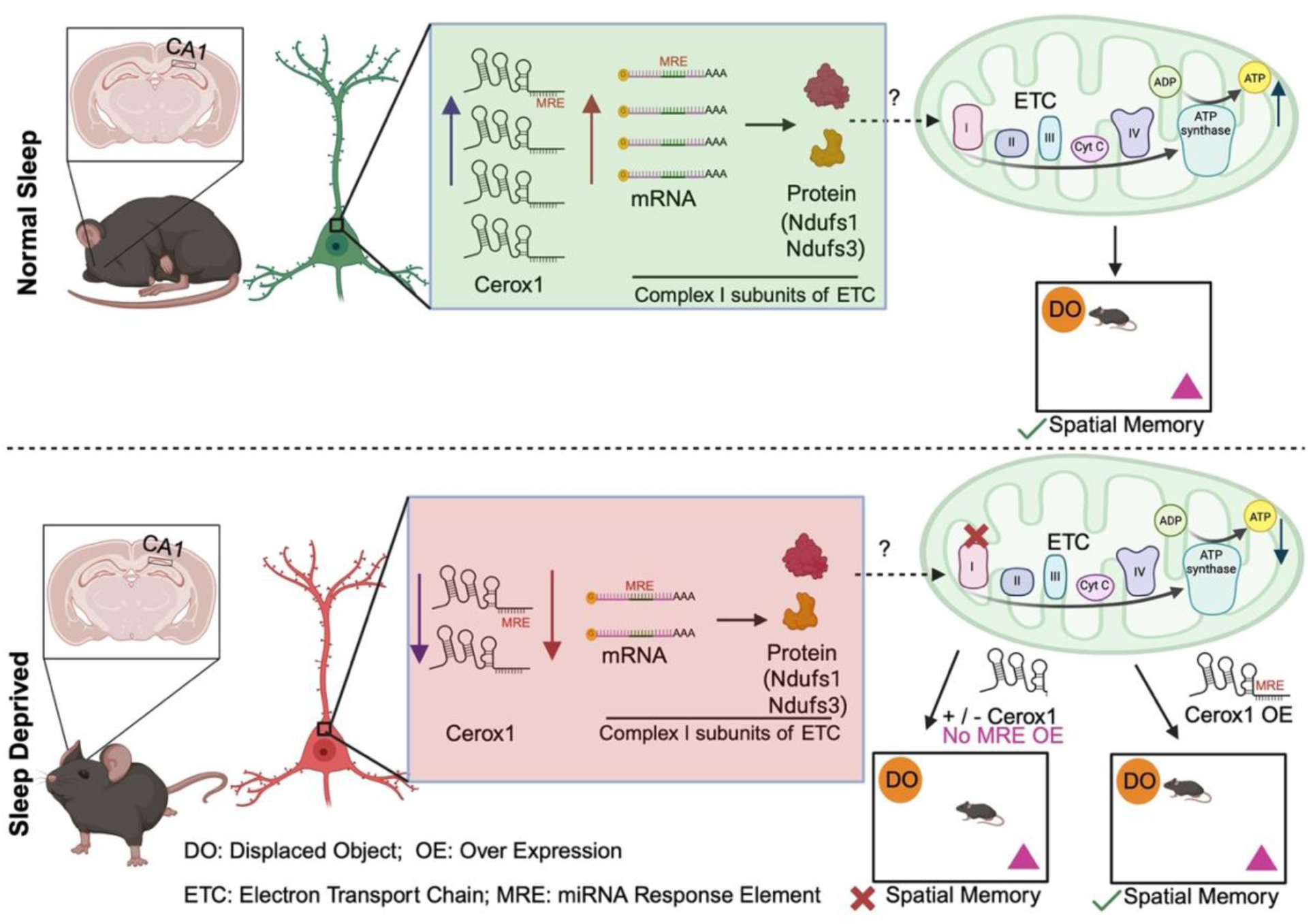

## Introduction

Sleep deprivation (SD) negatively affects cognitive function including memory storage [1–6]. A number of studies demonstrated that the hippocampus is especially susceptible to the detrimental effects of sleep loss, which disrupts key metabolic processes essential for memory storage [1–3, 7, 8]. Brief periods of SD for 5-6 hours have been shown to impair hippocampus-dependent memory consolidation [3, 7]. Among the various pathways regulated by sleep, energy metabolism has garnered increasing scientific interest [7]. Notably, SD has been reported to decrease the adenosine triphosphate (ATP) levels within the hippocampus [9]. This elevation of ATP levels is associated with decreased phosphorylation of adenosine monophosphate-activated protein kinase (pAMPK). pAMPK has been shown to function as cellular energy sensor by adjusting the ATP levels accordingly [7, 9]. For efficient use of limited energy resources after SD, the energy resources are redirected towards fundamental cellular process necessary for survival rather than being used for energy demanding processes, such as synaptic plasticity and memory. These observation suggests that this reallocation of energy recourses can potentially contribute to the memory impairments after SD [7]. However, the mechanistic understanding of how ATP synthesis is regulated after SD and how disruption of this process causes memory deficit are poorly understood.

Recent work has shown that the enhanced phosphorylation of AMPK negatively regulates protein synthesis in hippocampus after SD and disrupts memory consolidation [7, 10]. The deficit in protein synthesis via mTORC1 signalling cause impairment of hippocampus-dependent memory consolidation [10]. Given the established role of pAMPK as a cellular energy sensor that modulates protein synthesis in response to SD, it is reasonable to posit that sleep-dependent energy redistribution may be governed through translational regulatory mechanisms. Among the various molecular effectors of protein synthesis underlying memory formation, non-coding RNAs have been identified as pivotal regulators [11–15], including long non-coding RNAs or lncRNAs. These transcripts lack conserved coding potential but can function as regulators of diverse cellular processes including translation [12–15]. LncRNAs can function as a decoy to sequester translation repressor necessary for protein synthesis in neuronal dendrite and disruption of this decoy activity causes hippocampus-dependent memory deficit [13, 16]. Further, a recent study in non-neuronal system demonstrated that lncRNA could drive ATP synthesis by regulating the activity of mitochondrial electron transport chain (ETC). The overexpression of lncRNA promotes translation from transcripts encoding subunits of ETC and subsequently enhanced its activity [17]. While these findings suggest the involvement of RNA-mediated mechanisms in regulating cellular energy resources, the precise impact of sleep deprivation on energy availability and its contribution to memory impairment remains poorly understood.

In this study, we examined the role of lncRNA in memory deficit caused by sleep deprivation, a commonly observed symptom associated with diverse psychiatric conditions including stress, aging and neurodegeneration. We have identified Cerox1 as a sleep-regulated lncRNA that can rescue memory deficit after sleep deprivation. Our study uncovered the underlying mechanism of memory deficit that involves Cerox1 - mediated impairment of ETC activity necessary for ATP synthesis. We have shown that sleep deprivation or knockdown of Cerox1 reduced the ATP level in hippocampus, whereas the Cerox1 overexpression reversed it. We found that the overexpression of Cerox1 enhanced the expression of critical ETC complex I subunits, which share common microRNA response elements (MREs) with Cerox1. Furthermore, our data demonstrated that Cerox1 variant lacking MRE failed to rescue memory deficit caused by sleep deprivation. Collectively, our findings reveal a previously unrecognized mechanism of lncRNA-mediated ATP production and memory consolidation that is disrupted by sleep deprivation.

## Results

### Sleep regulates the expression of long non-coding RNAs

We focused on an emerging class of regulatory RNAs known as lncRNAs (>200 nucleotide long transcripts without conserved protein coding potential) [12, 14, 15]. Our recent transcriptomic analysis along with other studies identified a subset of lncRNAs in synaptosomes that regulate memory [16, 18, 19]. To pinpoint sleep-regulated lncRNAs, we have compared the set of synapse-enriched lncRNAs [16] with differentially expressed transcripts from the dorsal hippocampus following 5 hours of sleep deprivation (ZT0 to ZT5) [20]. We have measured biochemical changes from the dorsal hippocampus as the study focused on investigating the importance of these biochemical changes in the regulation of spatial memory after SD. This comparative assessment identified four lncRNAs that are downregulated after sleep loss. These include 2810468N07Rik (also known as <u>C</u>ytoplasmic <u>e</u>ndogenous <u>r</u>egulator of <u>ox</u>idative <u>p</u>hosphorylation 1 or Cerox1), Gm42770, AI504432 and Gas5 (Figure 1A). We further analyzed the differential expression of these lncRNAs in the hippocampus from mice (i) left undisturbed (NSD), (ii) after 5-hour sleep deprivation (SD), (iii) or after 2.5 hours of recovery sleep following sleep loss (RS). Efficacy of SD was assessed by the reduction in phosphorylation of translation initiation factor 4EBP2 as reported previously [10] (Figure 1B-1C). We observed that sleep deprivation reduced Cerox1 expression and recovery sleep reversed it (Figure 1D). Although Gm42770 showed significant recovery of expression after rescue sleep, the reduction of expression of this transcript after sleep deprivation is not statistically significant. Further, sleep deprivation or recovery sleep did not alter significantly the expression of AI504432 and Gas5 (Figure 1D). Based on the changes in expression, along with its evolutionary conservation and lack of protein coding potential [17] we chose to characterize the role of Cerox1 in memory impairment caused by sleep deprivation. To identify target transcripts of Cerox1 in the context of SD, transcriptomic data from Neuro2A cells overexpressing Cerox1 [17] was compared with differentially expressed transcripts in the hippocampus after sleep deprivation [20]. We observed an overlapping set of 90 transcripts in these two data sets. Gene Ontology (GO) analysis of these overlapping transcripts revealed a significant enrichment of GO-terms focused on oxidative phosphorylation, cellular respiration, and assembly of the mitochondrial electron transport chain (ETC) (Figure 1F), an observation consistent with the known impact of sleep loss on metabolism [7].

**Figure 1:**
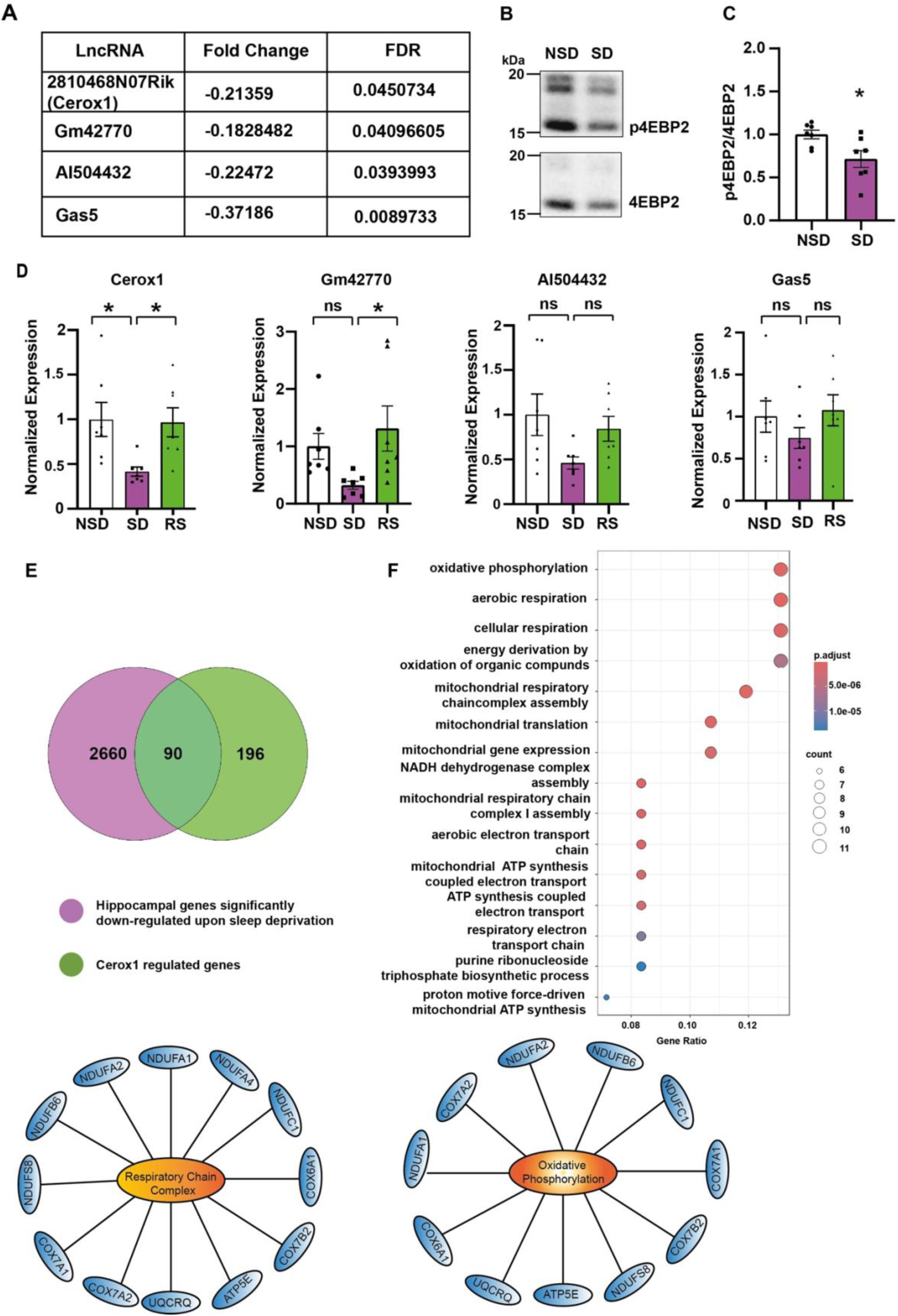
Identification of Cerox1 as a sleep-regulated lncRNA and its involvement in mitochondrial ATP synthesis. **(A)** Comparative analysis of differentially expressed, synapse-enriched lncRNAs from hippocampus identified four lncRNAs that are significantly down-regulated following sleep loss. **(B)** Western blot showing sleep deprivation leads to reduced phosphorylation of 4BEP2. **(C)** Quantification of Western blot in B. n= 7, *p = 0.0283, unpaired two-tailed *t-test* with Welch’s correction **(D)** qPCR analysis of Cerox1, Gm42770, AI504432 and Gas5 expression in the dorsal hippocampus after 5 hours of sleep deprivation (SD) or left undisturbed (NSD) or 2.5 hours of rescue sleep (SR) after SD. n = 7, Cerox1: F_2,18_ = 2.808, *p=0.035; Gm42770: F_2,18_ = 1.81, p=0.261; AI504432: F_2,18_ = 2.362, p=0.089; Gas5: F_2,18_ = 2.363, p=0.089, one-way ANOVA with Bonferroni’s correction **(E)** Comparative analysis of differentially expressed transcripts after SD and transcripts differentially expressed after Cerox1 overexpression identified 90 overlapping transcripts. **(F)** Gene Ontology (GO) analysis of the 90 overlapping transcripts showing enrichment of GO - terms involved in energy metabolism. ns = not significant. Data represented as mean ± SEM. “n” represents biological replicates.

### Sleep-regulated expression of Cerox1 modulates the activity of the mitochondrial electron transport chain

SD for 3 or 6 hours has been shown to prevent the increase in ATP that is commonly observed in distinct brain areas including the hippocampus during the initial hours of sleep [9]. We hypothesized that Cerox1 controls ATP levels during SD. Aerobic ATP synthesis is carried out by the mitochondrial ETC, which consists of five multi-subunit complexes [17]. The vast majority of ETC proteins are encoded by the nucleus; only 13 proteins are encoded by the mitochondrial genome [17]. Among the 90 genes regulated by both Cerox1 and SD, we focused on nuclear encoded ETC transcripts (shown in Figure 1F) as they show the highest increase in expression when Cerox1 is overexpressed. Consistent with these results, we observed that the sleep deprivation specifically reduced complex I activity without influencing complex II activity (Figure 2A).

**Figure 2:**
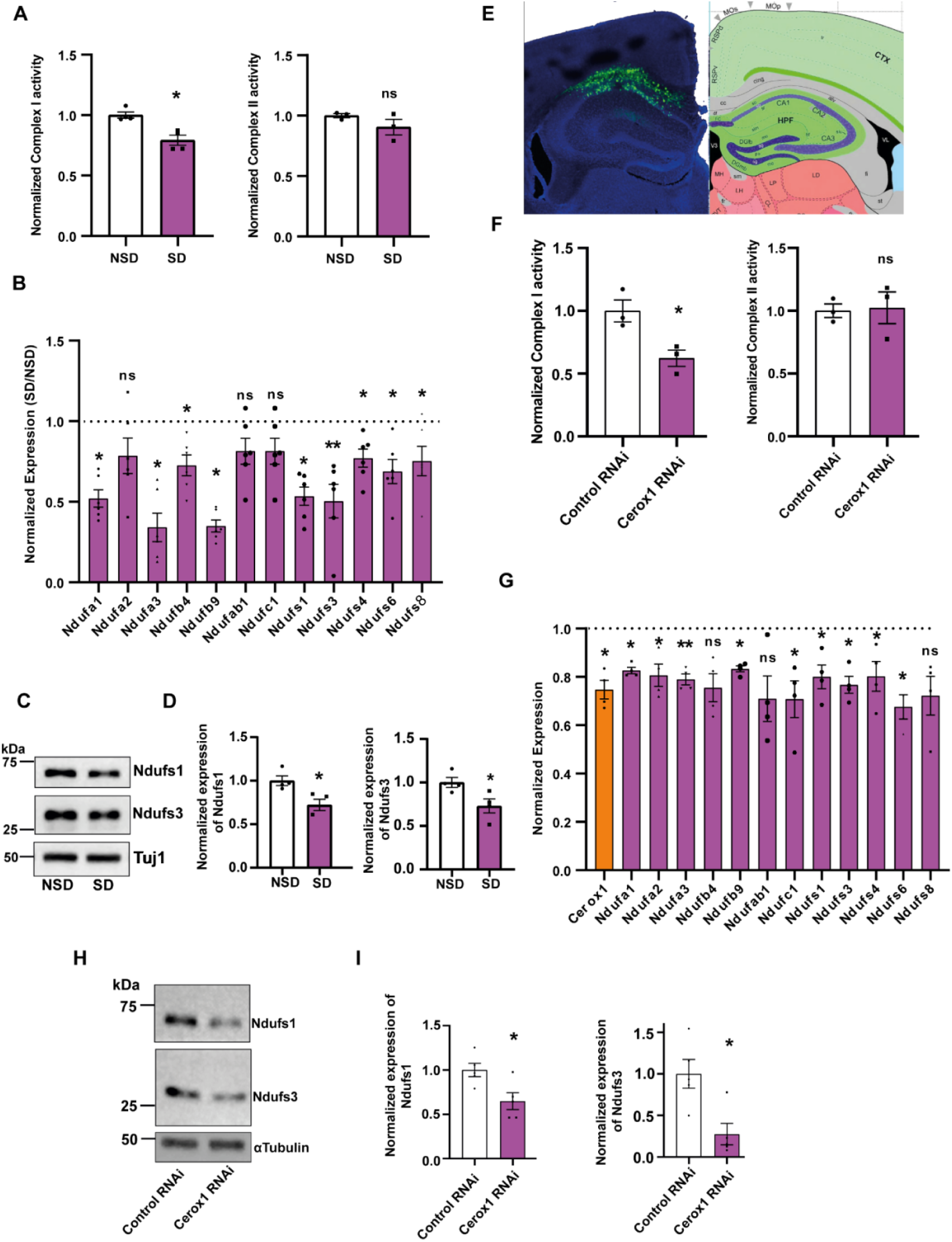
Sleep deprivation or Cerox1 knockdown negatively regulates mitochondrial electron transport chain activity. **(A).** SD reduces complex I activity without influencing complex II activity of ETC. Complex I activity, **p =0.0078; n = 4 for NSD or SD; complex II activity: n=3, p=0.27 unpaired two-tailed *t-test* with Welch’s correction. **(B).** qPCR profiling of 12 complex I subunits from the dorsal hippocampus from sleep deprived or undisturbed mice. n = 5, Ndufa1, *p =0.031; Ndufa2, p=0.159; Ndufa3, *p=0.024; Ndufb4, *p=0.013; Ndufb9, *p=0.021; Ndufab1, p=0.3; Ndufc1, p=0.129; Ndufs1, *p=0.028; Ndufs3, *p=0.007; Ndufs4, *p=0.05; Ndufs6, *p=0.017; Ndufs8, *p =0.048. Unpaired two-tailed *t-test* with Welch’s correction. **(C).** Western blot reduced expression of Ndufs1 and Ndufs3 from the dorsal hippocampus following SD or NSD. Tuj1 was used for normalization. (D) Quantitation of “C”. Ndufs1: n = 4, *p=0.018; Ndufs3: n = 4, *p=0.03; two-tailed *t-test* with Welch’s correction. **(E).** Dorsal hippocampus showing co-expression of eGFP along with Cerox1 or Control shRNA. **(F).** Cerox1 knockdown reduced Complex I, but not complex II, activity. n = 3, complex I: *p =0.02; complex II: p=0.878, unpaired two-tailed *t-test* with Welch’s correction. **(G).** qPCR profiling of 12 complex I subunits and Cerox1 from the dorsal hippocampus expressing Cerox1 shRNA or control shRNA. n = 4, Cerox1, *p=0.044; Ndufa1, *p =0.028; Ndufa2, *p=0.033; Ndufa3, **p=0.009; Ndufb4, p=0.083; Ndufb9, *p=0.023; Ndufab1, p=0.083; Ndufc1, *p=0.033; Ndufs1, *p=0.017; Ndufs3, *p=0.012; Ndufs4, *p=0.013; Ndufs6, *p=0.013; Ndufs8, p =0.107; unpaired two-tailed t-test with Welch’s correction. **(H).** Western blot showing reduced expression of Ndufs1 and Ndufs3 following knockdown of Cerox1. **(I).** Quantification of “H”. n = 5, Ndufs1: *p=0.021; Ndufs3: p=0.01. See Figure S1.

We validated our transcriptomic data of genes differentially regulated by SD and Cerox1 overexpression by qPCR. Indeed, SD led to significant reduction of transcripts encoding key proteins of complex I subunit. These include Ndfufa1, Ndufa3, Ndufb4, Ndufb9, Ndufs1, Ndufs3, Ndufs4, Ndufs6 and Ndufs8 (Figure 2B). However, the expression of Ndufa2, Ndufab1 and Ndufc1 remained unaltered. At the protein level, we observed that SD led to a significant reduction in Ndufs1 and Ndufs3 (Figure 2C – 2D).

To assess the impact of Cerox1 downregulation following SD, we first measured ETC activity upon Cerox1 knockdown by antisense RNA. Causally linking reduced Cerox1 expression to impaired complex I activity after SD, knockdown of Cerox1 in the dorsal hippocampus reduced complex I, but not complex II, activity (Figure 2E – 2F). Further, the knockdown of Cerox1 either *in vivo* or *in vitro* significantly reduced the expression of Ndufa3, Ndufs1, Ndufs3 and Ndufs6 transcripts (Figure 2G and S2).

At the protein level, the expression of Ndufs1 and Ndufs3 was reduced after knockdown of Cerox1 (Figure 2H – 2I). To analyze the specificity of Cerox1-dependent expression of complex I subunit transcripts after sleep deprivation, we measured the levels of another transcript that is differentially regulated by SD, Cold Inducible RNA-Binding Protein (CIRBP). Consistent with previous studies [21, 22], the expression of CIRBP was reduced after sleep deprivation (Figure S1C – S1D) but its expression did not change after Cerox1 knockdown (Figure S1E – S1F).

Having shown that Cerox1 knockdown phenocopies mitochondrial dysfunction after SD, we asked whether Cerox1 overexpression might have the opposite effect. Indeed, adenovirus-mediated overexpression of Cerox1 enhanced complex I activity without affecting complex II function (Figure 3A). Consistent with these data, the expression of Ndufa3, Ndufs1, Ndufs3 and Ndufs6 transcripts, all subunits of complex I, were enhanced after Cerox1 overexpression (Figure 3B). At the protein level, the expression of Ndufs1 and Ndufs3 was enhanced after Cerox1 overexpression (Figure 3C – 3D). We have measured the ATP level from the dorsal hippocampus of SD or NSD mice after Cerox1 or tdTomato overexpression. Consistent with previous observation [9], SD led to a significant reduction of ATP and this reduction was partially reversed after Cerox1 overexpression (Figure 1E). We evaluated the expression of Hsp60, a constitutively expressed mitochondrial matrix protein, as a proxy for mitochondrial mass. We observed that SD did not alter the levels of mitochondrial Hsp60 relative to cytosolic β-tubulin (Figure 3F – 3G).

**Figure 3:**
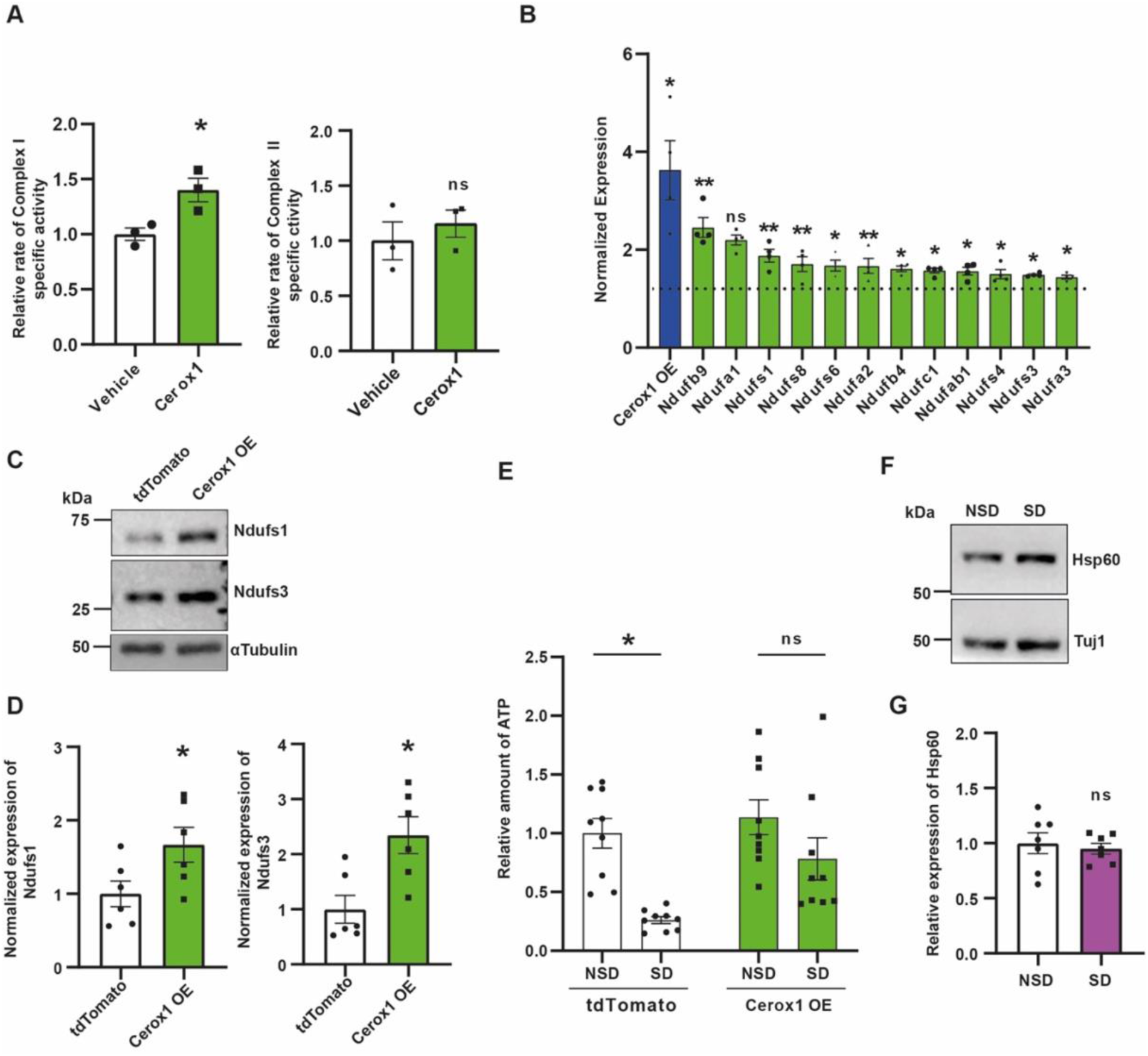
Cerox1 overexpression enhances mitochondrial electron transport chain activity and reverses sleep deprivation -induced decrease in ATP **(A)** Cerox1 overexpression (OE) enhances complex I activity without influencing complex II activity of ETC. Complex I activity: *p =0.043; n=3 for tdTomato or Cerox1 OE, complex II activity, p=0.504 unpaired two-tailed *t-test* with Welch’s correction. **(B).** qPCR profiling of 12 complex I subunits and Cerox1 from the dorsal hippocampus overexpressing Cerox1 or tdTomato. n = 4, Cerox1, *p=0.014; Ndufa1, **p =0.002; Ndufa2, p=0.056; Ndufa3, **p=0.002; Ndufb4, **p=0.008; Ndufb9, *p=0.044; Ndufab1, **p=0.004; Ndufc1, *p=0.021; Ndufs1, *p=0.041; Ndufs3, *p=0.028; Ndufs4, *p=0.016; Ndufs6, *p=0.02; Ndufs8, *p =0.013. unpaired two-tailed t-test with Welch’s correction. **(C).** Western blot showing expression of Ndufs1 and Ndufs3 after overexpression of Cerox1 or tdTomato. αTubulin used for normalization **(D).** Quantification of western blot. n = 6, Ndufs1: *p=0.048, Ndufs3: *p=0.01, unpaired two-tailed *t-test* with Welch’s correction. **(E).** Luminescence based quantitation of ATP from SD and NSD mice after Cerox1 or tdTomato expression. n = 9, Sleep deprivation: F_1,32_ = 16.76, **p= 0.0003; Cerox1 OE: F_1,32_ = 6.065, *p= 0.0194; two-way ANOVA with Tukey’s multiple comparison tests **(F).** Western blot showing expression of Hsp60, a mitochondria-localized protein chaperone, in SD and NSD mice. **(G).** Quantification of western blot. n = 6, p=0.65, unpaired two-tailed *t-test* with Welch’s correction. ns = not significant. Data represents mean ± SEM. “n” represents biological replicates. See Figure S2.

### Cerox1 delivery prevents memory impairment caused by sleep deprivation

SD for 5 hours causes hippocampus-dependent memory deficits [1, 3, 4, 10]. Sleep loss or knockdown of Cerox1 inhibits mitochondrial complex I activity. In contrast, the overexpression of Cerox1 enhanced complex I activity. Previous studies have shown that sleep deprivation reduces levels of ATP [9] and impairs memory [1, 3, 4]. Consistent with a previous study [9], we observed a reduction in ATP after SD. Importantly, this reduction in ATP levels was prevented when mice overexpressing Cerox1 were challenged by SD. Prompted by these observations, we investigated whether restoring Cerox1 levels in the dorsal hippocampus of sleep deprived mice would prevent memory deficits.

We used a spatial object recognition (SOR) task to measure hippocampus-dependent memory following exposure to SD [10, 23]. AAVs overexpressing Cerox1 or tdTomato were injected into the dorsal hippocampus and these mice were trained for the SOR task. Mice were allowed to explore and learn the position of two distinct objects within an arena. The spatial task exploits the preference for a displaced object as an indicator of memory. Immediately after training, mice overexpressing Cerox1 or tdTomato were sleep deprived for 5 hours or left undisturbed. The next day these mice were tested for objection-recognition memory (Figure 4A). In agreement with previous studies [3, 4, 10], mice expressing tdTomato that were left undisturbed showed enhanced preference for the displaced object, indicating consolidated memory (Figure 4B – 4C). Sleep deprived mice expressing tdTomato did not show enhanced preference for the displaced object, demonstrating a deficit in memory consolidation caused by sleep loss. Mice overexpressing Cerox1 showed enhanced preference for the displaced object, regardless of whether they were sleep-deprived or not (Figure 4B – 4C). This enhanced preference for the displaced object is comparable to tdTomato expressing mice that were left undisturbed. These observations indicate that Cerox1 overexpression is sufficient to rescue the impairment of hippocampus-dependent memory consolidation caused by sleep deprivation. Further, Cerox1 overexpression in absence of sleep deprivation showed enhancement in memory, but this increase is not statistically significant (Figure 4C). This observation prompted us to speculate that the complete rescue of SD -induced memory deficit, but not ATP level, may occur *via* Cerox1 -dependent other pathways in addition to regulation of mitochondrial ETC activity. Cerox1 transcripts contains miRNA response elements (MRE) and the overexpression of Cerox1 with MREs promotes translation from transcripts encoding subunits of the ETC including Ndufs1 and Ndufs3 [17]. In contrast, mutating the MREs was sufficient to block translation of these ETC subunits. We investigated whether Cerox1 overexpression without the MREs would affect memory consolidation after SD. Our findings showed that SD reduced exploration of the displaced object, and this diminished preference remained unchanged when Cerox1 lacking the MRE was overexpressed (Figure 4D–4E).

**Figure 4:**
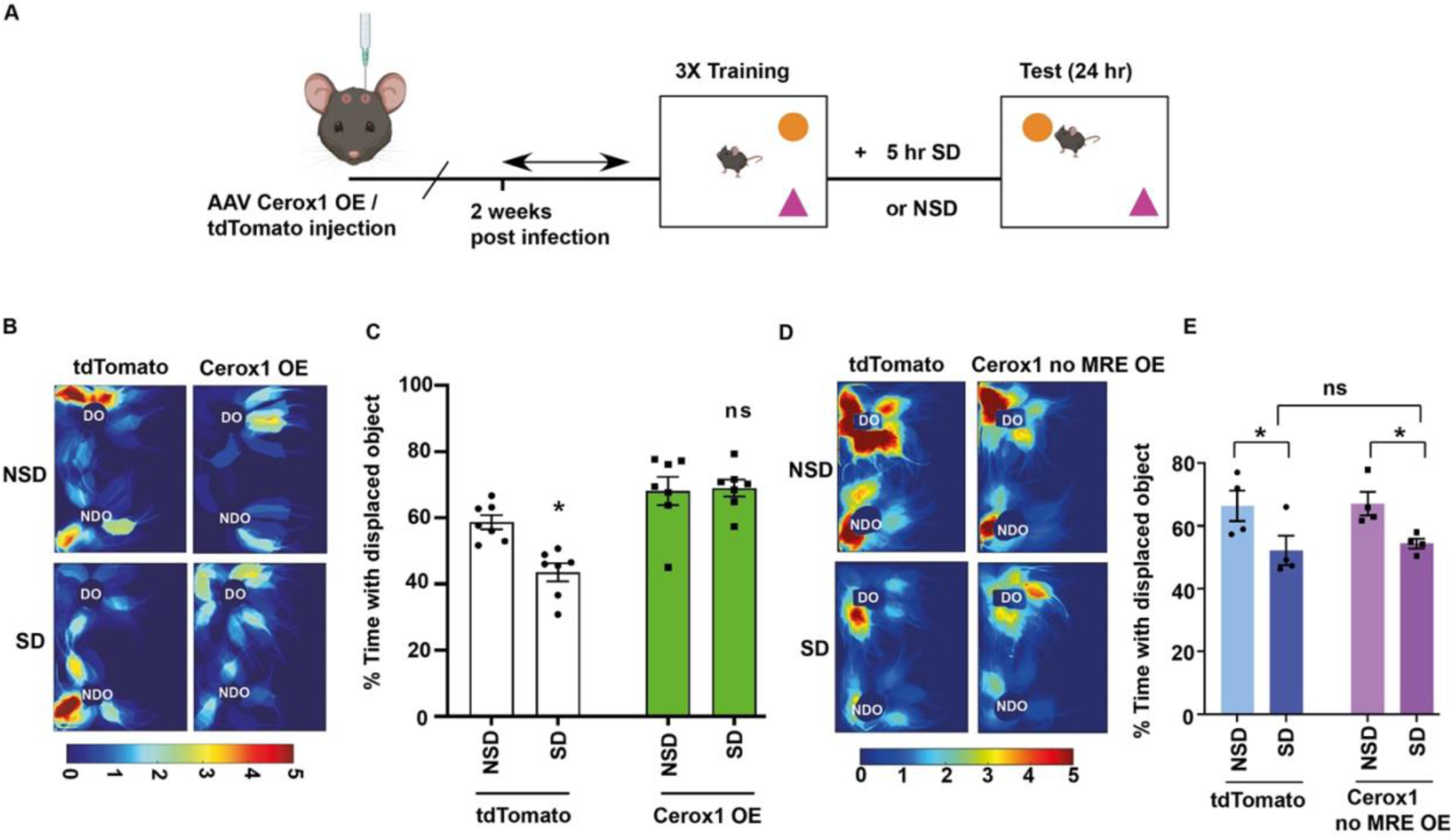
Cerox1 overexpression prevents memory impairments caused by sleep deprivation **(A).** Schematic representation of hippocampus-dependent spatial object recognition (SOR) test following sleep deprivation of mice overexpressing Cerox1 or tdTomato. Mice were trained for spatial object recognition prior to 5 hours of sleep deprivation or left undisturbed. Preference for the displaced or the non-displaced object were assessed 24 hours after SD. **(B).** Representative heat map displays mice expressing tdTomato showing reduction in the preference for exploring the displaced object after SD, whereas mice overexpressing Cerox1 showed increase in preference for the displaced object even after SD. Scale as indicated. **(C).** Quantification of preference for the displaced object represented as % time with displaced object. n = 7, sleep deprivation: F_1,24_ = 5.464, p = 0.0281; Cerox1 OE: F_1,24_ = 32.48, **p=0.0001; Interaction effect: F_1,24_ = 6.862, p=0.015; two-way ANOVA with Bonferroni’s corrections. **(D).** Representative heat map showing reduction in the preference for exploring displaced object after SD and this reduced preference for the displaced object after SD did not change when Cerox1 lacking MRE was overexpressed. Scale as indicated. **(E).** Quantification of preference for the displaced object represented as % time with displaced object. n = 4, sleep deprivation: F _(1,12)_ = 11.78, p = 0.005; Cerox1 no MRE: F _(1,12)_ 0.1385, p = 0.7163; Interaction effect: F _(1,12)_ = 0.03227, p = 0.8604; two-way ANOVA with Fischer’s LSD. Data represents mean ± SEM. “n” represents biological replicates. ns = not significant. MRE: MiRNA Response Element.

## Discussion

Emerging studies have demonstrated that lncRNAs, a class of regulatory non-coding RNA, are pivotal regulators of memory [16, 19]. However, the role of lncRNAs in memory deficits caused by sleep deprivation and mechanistic details of underlying processes are unknown. This study identifies Cerox1 as a sleep-regulated lncRNA (Figure 1). We show that memory deficit caused by sleep deprivation is mediated by Cerox1-dependent control of mitochondrial ETC activity necessary for ATP production (Figures 2, 3). We demonstrate that SD or knockdown of Cerox1 reduces the activity of ETC leading to decline in ATP, whereas overexpression of the lncRNA has an opposite effect (Figures 2, 3). Further, the overexpression of Cerox1 corrects memory deficits caused by SD (Figures 4).

How does Cerox1 regulate the ETC activity? Cerox1 is a nuclear encoded transcript that is predominantly localized in the cytoplasm and is not present within the mitochondria (Figure S1C). The nuclear-encoded subunits of the ETC are translated in the cytoplasm and then imported into the mitochondria, where they assembled into complexes with the mitochondrially encoded subunits [24–26]. The ETC is composed of five complexes. Complex I through IV set up the electrochemical gradient that complex V uses to synthesize ATP. Complex I contains the Core-Q module, Core N module and 31 supernumerary subunits. The Core-Q module is essential for electron transfer to ubiquinone, the Core-N module is required for NADH oxidation, and the supernumerary subunits assists the catalytic function of the complex without directly participating in catalysis [27]. We measured protein levels of Ndufs1 and Ndufs3 after altering the Cerox1 expression or SD because these two subunits are critical for catalytic activity of Core-N and Core-Q modules respectively. Furthermore, our results establish the specificity of Cerox1 in modulating the expression of Ndufs1 and Ndufs3 following SD, as its knockdown had no effect on the expression of CIRBP, a transcript also regulated by sleep (Figure S1C-1F). A study by Sirey et al. showed that the Cerox1 and these subunits of complex I share a common miRNA response element (MRE) [17]. Comparative analysis of this study and our data indicate that Cerox1 could regulate the expression of these subunits *via* sleep-regulated binding of miRNAs to these MREs. It is plausible that sleep facilitates the interaction between miRNAs and the Cerox1 lncRNA, thereby sequestering miRNAs that would otherwise repress the translation of transcripts encoding complex I subunits of the ETC. This hypothesis is supported by our data showing that overexpression of Cerox1 increases, while SD or Cerox1 knockdown reduces, the protein levels of Ndufs1 and Ndufs3—both essential components of complex I (Figure 2 - 3). In alignment with these findings, both SD and Cerox1 knockdown diminished complex I activity, whereas Cerox1 overexpression restored it (Figures 2–3). Moreover, SD resulted in reduced ATP synthesis, which was reversed by overexpressing Cerox1 containing the MRE (Figure 4). We observed that the decline in ATP levels caused by SD is greater than the reduction in complex I activity after Cerox1 knockdown. Considering prior evidence that Cerox1 regulates complex IV activity [17], we propose that Cerox1-mediated regulation of ATP synthesis during sleep may also involve complex IV activity. In agreement with our results, Sirey et. al. reported that overexpression of MRE-containing Cerox1 enhances ATP production [17]. They proposed that miRNAs are redistributed from Cerox1 to transcripts encoding complex I subunits, thereby promoting their translation, enhancing ETC function, and increasing ATP levels [17]. Sleep deprivation may disrupt this redistribution process, leading to impaired ETC activity and decreased ATP production. Furthermore, we demonstrated that overexpression of Cerox1 with an intact MRE rescued the memory deficits caused by SD, whereas the MRE-deficient variant failed to do so (Figure 4). Together, our findings underscore the physiological importance of miRNA–lncRNA interactions in regulating memory consolidation, a mechanism that is disrupted by sleep loss (Figure 4).

The energy hypothesis of sleep proposes that sleep pressure develops due to the depletion of cellular energy during wakefulness [7, 28]. Sleep deprivation reduces levels of ATP, disrupting the energy balance [7, 9]. Another set of studies showed that SD causes memory deficits [3, 7, 10]. However, the link between decline in ATP and memory deficit after sleep deprivation is not yet established. Our findings provide evidence linking these two sleep-regulated processes (Figures 3 - 4). A previous study [9] also reported a similar decline in ATP levels after 3 or 6 hours of sleep deprivation. This previous research demonstrated that the reduction in ATP is directly associated with sleep deprivation, rather than being influenced by circadian rhythm factors. Providing a molecular explanation for these findings, our data revealed that the decline in ATP levels after sleep deprivation depends on Cerox1 (Figure 3), while Cerox1 expression remains constant throughout the circadian cycle (Figure S1A-1B). Mechanistically the decline in ATP and impairment of late-phase long-term potentiation after sleep deprivation is regulated by AMPK signalling [9, 29–31]. Furthermore, AMPK signalling regulates protein synthesis [10]. Among various regulators of protein synthesis, lncRNAs emerged as a pivotal player [13, 16, 19]. We anticipate that future experiments will establish the direct link between lncRNA and AMPK signalling in the context of energy balance and memory.

Why does the reduction in ATP levels during sleep deprivation cause memory deficits? Sleep loss appears to redirect energy resources away from less immediately critical processes, like synaptic plasticity [7], to prioritize the minimal energy required for sustaining basic cellular physiology. Our findings demonstrate that the ectopic expression of Cerox1 during sleep deprivation counteracts this energy imbalance by restoring ATP levels (Figure 4). This restoration of ATP level rescues memory deficit caused by sleep loss. In contrast to our data and observation from Dworak et al. [9], another study by Natsubori et al. reported a decline in ATP during rapid eye movement (REM) sleep [32]. This difference may be due to the varying methods used to measure ATP levels. Natsubori et. al. employed a genetically encoded ATP sensor that specifically measures intracellular ATP [32], whereas this study and Dworak et al. used a luminescence-based assay to evaluate ATP from cellular lysates [9].

Sleep loss is a common symptom in neurodegenerative diseases like Alzheimer’s and Parkinson’s [33–35] and neurodevelopmental disorders, such as autism [36, 37]. Moreover, a severe impairment of the complex I activity [35, 38] is also reported in neurodegenerative disorders. Emerging evidence from large-scale genetic association studies implicates mitochondrial metabolism in the pathophysiology of psychiatric disorders [39–42]. Our work is the first to link sleep and memory with the RNA-mediated regulation of energy metabolism, laying the groundwork for future investigations that explore the potential of emerging RNA-based therapeutics.

## Acknowledgement

We thank Michael Kiebler, Max Harner and Sarbani Samaddar for helpful discussions and insightful comments. Balakumar Srinivasan for manuscript preparation.

## Funding

This work is supported by Department of Biotechnology (BT/PR45525/MED/122/311/2022) and core fund from National Brain Research Centre. TA is the Roy J. Carver Director of the Iowa Neuroscience Institute and is supported by NIH R01 AG 062398. SS is supported by NIH R21 AG080472-01, the Ataxia Charlevoix-Saguenay Foundation, Regents of the University of California, A22-2853-S004 (Jordan’s Guardian Angels), DoD/CDMRP (AR230307), the Eagles Autism Foundation, and the Simons Foundation Autism Research Initiative (SFARI).

## Conflict of Interest

All authors declared that there is no conflict of interest.

---

**Figure S1:**
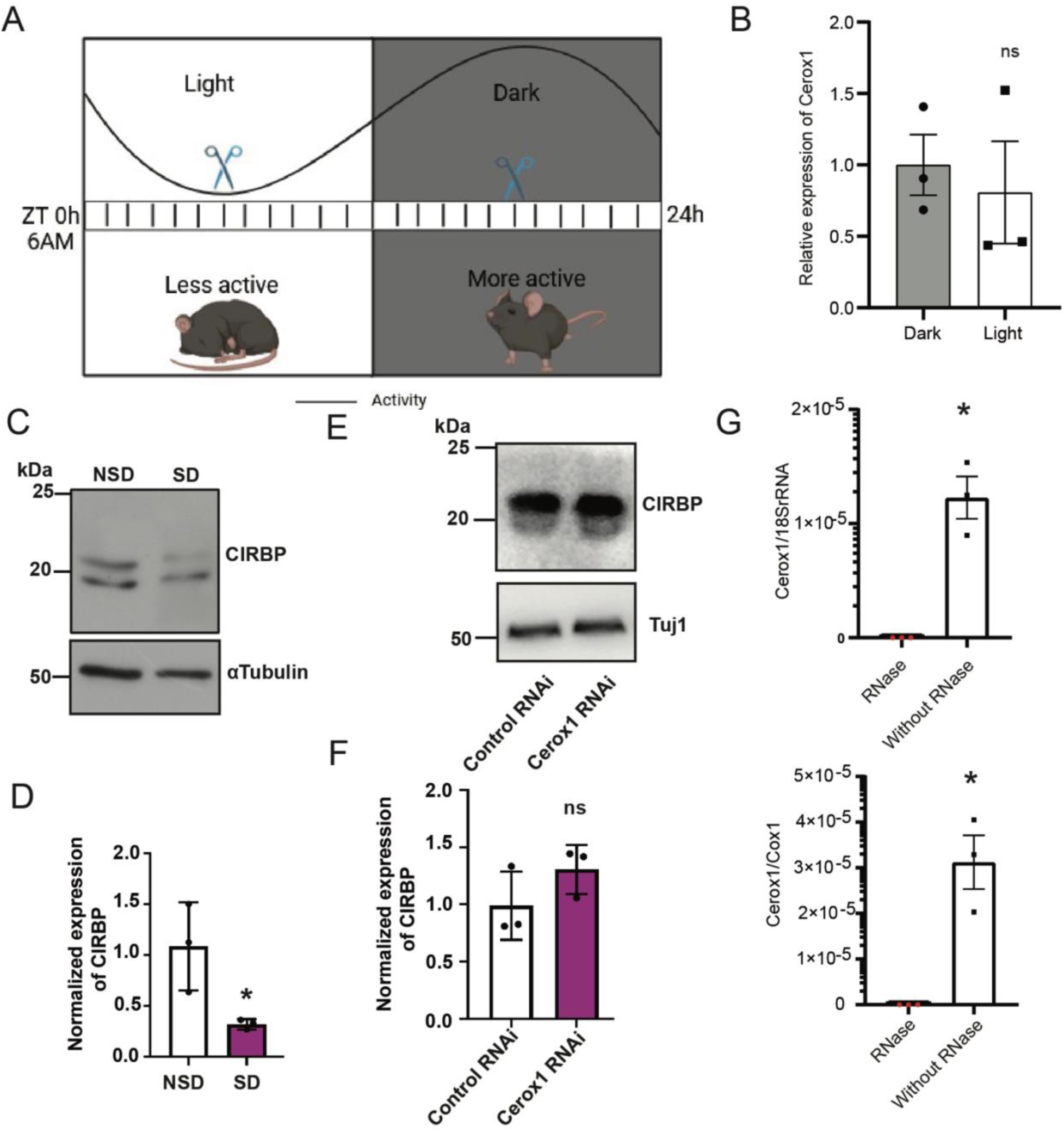
Abundance of Cerox1 during circadian cycle and in mitochondria **(A).** Schematic representation of circadian cycle showing sleep and awake cycle. Hippocampus was harvested as indicated during the cycle. **(B).** qPCR analysis of Cerox1 abundance in Dark (awake) and Light (sleep) phase of circadian cycle. **(C).** Western blot showing expression of CIRBP from NSD or SD mice. **(D).** Quantification of western blot. n = 3, *p = 0.0383, unpaired two tailed *t-test*. **(E).** Western blot showing CIRBP expression in presence or absence of Cerox1. **(F).** Quantification of western blot. n = 3, unpaired two tailed *t-test*. ns = not significant. **(G).** Abundance of Cerox1 in mitochondrial fraction treated with or without RNase. qPCR analysis showing Cerox1 expression as normalized by 18S rRNA and mitochondria specific Cox1 transcript. n = 3, Data represents mean ± SEM. “n” represents biological replicates. Data represents mean ± SEM. “n” represents biological replicates.

**Figure S2:**
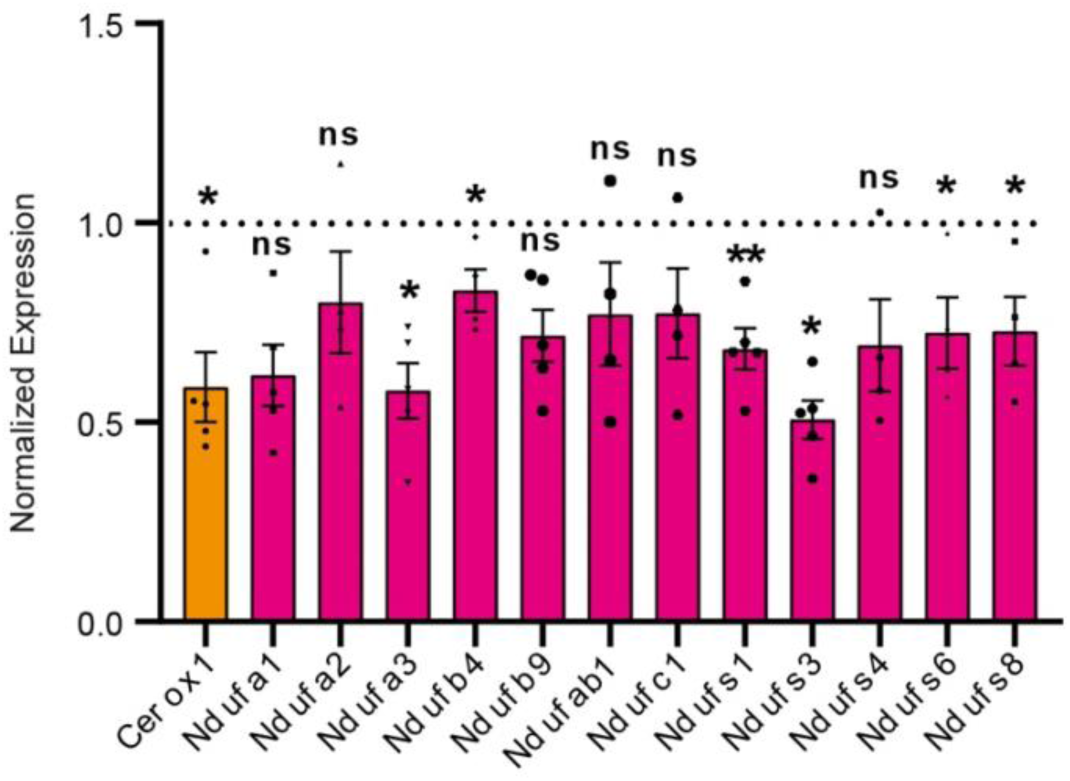
Expression of complex I subunits following Cerox1 knockdown in primary hippocampal neurons. qPCR profiling of 12 complex I subunits and Cerox1 from the dorsal hippocampus expressing Cerox1 or control shRNA. Cerox1, n=5, *p=0.047; Ndufa1, n=5, p =0.056; Ndufa2, n=4, *p=0.219; Ndufa3, n=5, *p=0.026; Ndufb4, n=4, *p=0.042; Ndufb9, n=5, p=0.07; Ndufab1, n=4, p=0.174; Ndufc1, n=4, p=0.088; Ndufs1, n=5, **p=0.0016; Ndufs3, n=5, *p=0.034; Ndufs4, n=4, p=0.07; Ndufs6, n=4, *p=0.046; Ndufs8, n=4, *p =0.032. Statistical analysis between groups performed using two-tailed t-test with Welch’s correction. Data represents mean ± SEM. “n” represents biological replicates.

## Methods

### Sleep deprivation

All animal experiments in this study were conducted with the approval of Institutional Animal Ethics and Care committee (NBRC/IAEC/2024/193 and NBRC/IAEC/2020/169). For sleep deprivation studies male C57BL/6J mice (Jax, #000664) of 8-12 weeks age were divided into Non-Sleep Deprived (NSD), Sleep Deprived (SD) and Recovery Sleep (RS) groups and housed in individual cages with corncob bedding on a 12 h light/dark cycle with lights ON at 06:00 AM (ZT0). Water and food supplied in *ad libitum* during the entire course of experiment. Animals were habituated for 5 mins each for 3-6 days with gentle handling techniques as described in previous studies (1, 2). On the day of the experiment animals grouped under SD and SR group were deprived of total sleep for 5 hours using the gentle handling techniques starting from ZT0. SD mice were subjected to recovery sleep for 2.5 hours (RS). Following sleep deprivation, NSD and SD animals were euthanized by cervical dislocation, and their bilateral dorsal hippocampal region was isolated immediately. Subsequently, the dorsal hippocampus was isolated from SR animals. All dissected dorsal hippocampal regions were immediately homogenized in RIPA (Millipore, #20 -188) supplemented with protease inhibitor (Sigma, #P8340) and phosphatase inhibitor (Sigma, #P0044) or TRIzol™ (Invitrogen, #15596026) and processed for biochemical experiments.

### Quantitative PCR

Total RNA isolated from the dorsal hippocampus using TRIzol^TM^ reagent as directed by manufacturer’s instruction. The tissues were homogenized by passing through needle and isolated through phase extraction using chloroform. The RNA was precipitated using isopropanol and washed in the ethanol. The isolated total RNA then quantified using Nanodrop (Thermo Scientific™ 2000/2000c Spectrophotometer) and their integrity was verified using an RNA gel. 2 µg of the total RNA was used for cDNA synthesis with random hexamers using Superscript III (Invitrogen, # 18080051) after removal DNase treatment of total RNA (Invitrogen, #AM1906). The abundance of transcripts were quantified by qPCR using gene specific primers (See Table 1). The qPCR was performed using Power SYBR Green (Applied Biosystems^TM^, #4367659) as instructed by the manufacturer’s in CFX Opus 96 Real-Time PCR System (Bio-Rad, #12011319). The relative expression of the transcripts was measured using ΔΔC_t_ method. The expression was normalized by the the β-tubulin expression.

**Table 1.**
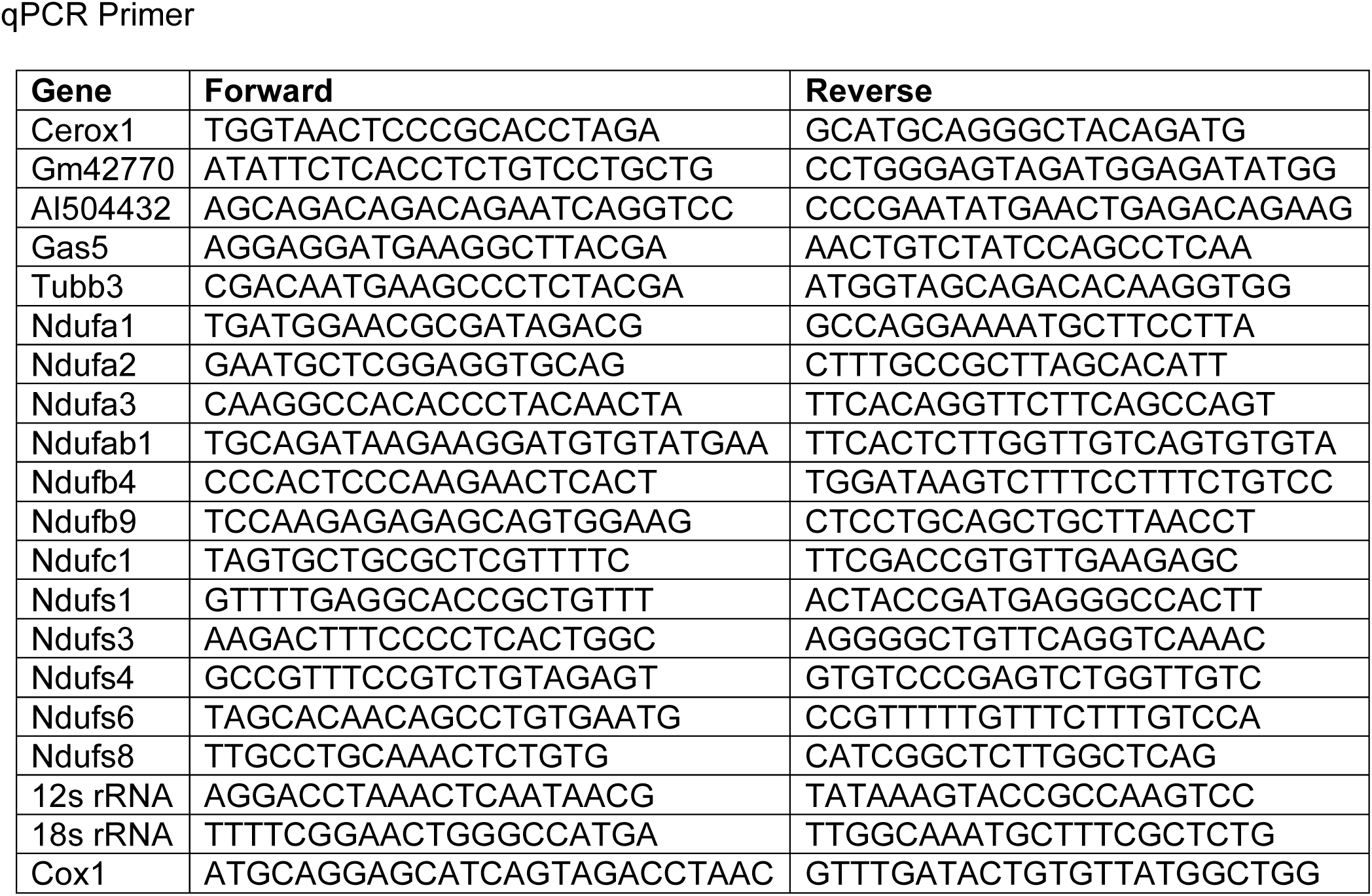
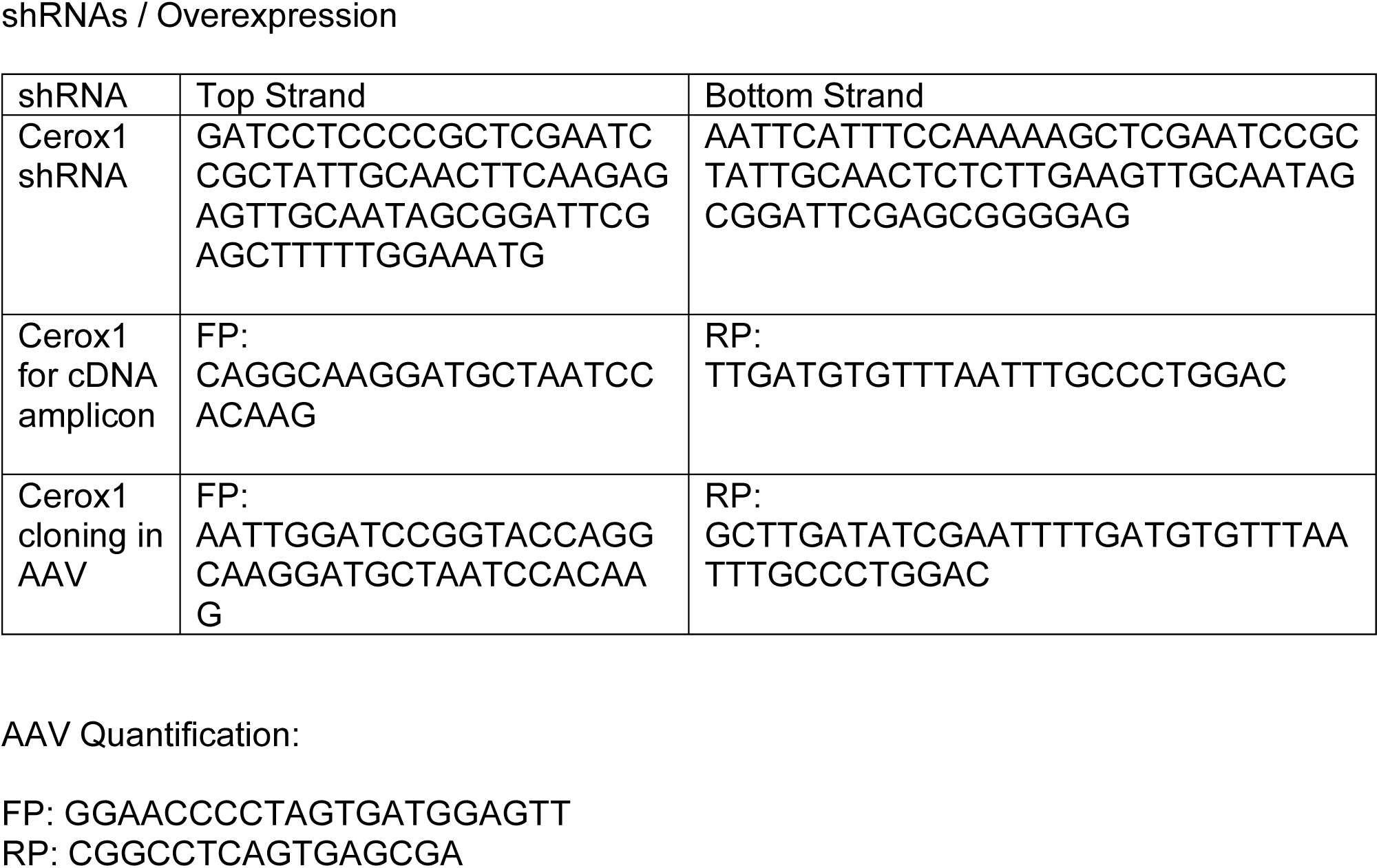
qPCR Primer

### Western blotting

Protein concentration of the dorsal hippocampal lysate was quantified using the BCA method (Pierce, #23227) as directed by the manufacturer. The samples were then electrophoresed using 10% or 15% sodium dodecyl sulfate-polyacrylamide gel (SDS-PAGE) in Tris-glycine buffer and transferred on to nitrocellulose membrane. The membrane was then blocked using 5% bovine serum albumin (BSA) in tris buffered saline (pH7.4) containing 0.1% Tween 20 or 5% skimmed milk in phosphate buffered saline (pH7.4) containing 0.1% Tween 20 (PBST) at room temperature for 1 hour. The blot was then incubated with primary antibody against the protein of interest in the blocking solution overnight at 4°C on a rocker. The primary antibody for probing of phosphorylation-sensitive p4EBP1 (Cell Signaling Technology, #2855; detect p4EBP2 as it is predominantly present in brain) diluted at 1:1,000 4EBP2 (Cell signaling Technology, #2845) diluted at 1:2500 and CIRBP (Abcam, #ab106230) diluted at 1:1000 using BSA in TBST. The antibodies used are as follows: Ndufs1 (Abcam, #ab169540) at 1:30,000 dilution, Ndufs3 (Abcam, #ab110246) at 1:3,000, Hsp60 (Cell signaling, #12165S) at 1:2000 dilution and houskeeping control genes, β-tubulin III (Sigma-Aldrich, #8578) at 1:30,000 and α-Tubulin (Sigma-Aldrich, #T5168) at 1:25,000 dilutions membrane were blocked with 5% skimmed milk in PBST. Membrane were then washed and probed with HRP-tagged secondary antibodies. Membranes were then washed and imaged with Immobilon signal enhancer (Millipore, #WBKLS0500), using UVI-Tech system. The chemiluminescent signal density was quantified using Bio-Rad Image lab software tool (version 6.0.1 standard edition). The density of p4EBP2 normalized to the expression of 4EBP2 and others were normalized to the signal density of β-tubulin III or α-tubulin.

### Analysis of Cerox1-Regulated Genes

We examined the effect of sleep deprivation on Cerox1 -regulated genes by analyzing a set of 286 genes, comparing their expression changes upon sleep deprivation. To identify potential biological implications, we performed enrichment and pathway analysis using the KEGG database. Specifically, we focused on the 90 Cerox1-regulated genes that were significantly downregulated upon sleep deprivation. This analysis was carried out using the cluster profiler (Version 4.14.4) package in R (Version 4.4.2) / Bioconductor (Version 3.19) (3). The results were then visualized as bubble plot using enrichplot package (Version 1.26.3)

### Cloning of knock-down and overexpression constructs

The knockdown construct generated by cloning short hairpin RNA (shRNA) cassette (See Table 1) targeting Cerox1 into pAAV.U6.shRLuc.CMV.zsGreen.SV40 plasmid (Addgene # 163020) under the U6 promoter. The shRNA cassettes were chemically synthesized (Merck oligos) with overhang for BamHI and EcoRI site to enable directional insertion into the target vector. The recombinant product then sequence verified using Sanger sequencing and used for the AAV preparation. For the overexpression construct the endogenous Cerox1 gene along with its predicted MRE element was PCR cloned from mouse hippocampus derived cDNA. The amplified gene products were incorporated into the pDrive vector using TA cloning (Qiagen, #231124). The positive clones were then chosen using blue-white screening and sequencing was verified with DNA-Sanger sequencing. The AAV Cerox1 construct was then made by infusion cloning (Takara Bio, #638947) of the verified Cerox1 with MRE sequence (∼1.2 kbp) into AAV-CAG-tdTomato (Addgene plasmid # 59462; RRID:Addgene_59462) replacing the tdTomato sequence under the chicken β-actin promoter. The resultant plasmid was then verified and used for the AAV-viral preparation.

### AAV vector preparation

HEK 293T cells maintained in Dulbecco’s modified eagle medium (Gibco, # 31600083) supplemented with 10% Fetal Bovine Serum (Gibco, # 10082147) and penicillin streptomycin (Gibco, #15140122) cocktail used for the production of replication incompetent recombinant AAV viral vectors. The HEK293T cells co-transfected at a ratio of 2:1:1 with plasmids pAAV2/9n (Addgene plasmid #112865; RRID: Addgene_112865), pAdDeltaF6 (Addgene plasmid #112867; RRID: Addgene_112867) and the recombinant plasmids carrying knockdown or Cerox1 overexpression constructs or their respective controls using calcium phosphate transfection method. Following three days of incubation the AAV virus released into the contact medium and bound to the cells isolated by freeze thaw cycles and concentrated using ultracentrifugation (Beckman coulter, Optima L100) with sucrose cushion. The AAV viral pellets was then re-suspended in PBS containing 5% (v/v) of glycerol and stored in aliquots at -80°C freezers. The AAV viral genome copy (GC) was then quantified using quantitative PCR (See Table 1). The viral titration was in the range of 1.5 x 10^13^ GC/mL to 3.7 x 10^13^ GC/mL. Around 1 µL of titer adjusted AAV viral suspension was used in the stereotaxic injection delivery and for the transduction of cultured hippocampal neurons.

### Primary hippocampal neuronal culture preparation

The conditioned media from astroglial feeder cells were prepared first to support the growth and maintenance of the primary hippocampal culture. Newborn CD1 pups were euthanized by carbon-di-oxide chambers and their brains were isolated and transferred into dissection media made of Hank’s Balanced Salt Solution (Sigma-Aldrich, #H2387) buffered with 10 mM HEPES (free of calcium, magnesium and bicarbonate) in aseptic hood. Using dissection microscope the meninges were carefully removed and the hemispheres are minced with surgical blade. Following that tissues were dissociated with 2.5% Trypsin (Gibco, #15090046) and 1% (w/v) DNase (Sigma-Aldrich, # DN25). The cell suspension then filtered with nylon filter to eliminate undissociated chunks. The filtrate then centrifuged at 120 x g and the pellet re-suspended and plated in MEM medium (Gibco, #11095080) supplemented with 20% (w/v) glucose (Gibco, # 15023021), 100X pyruvate (Gibco, #11360-070), horse serum (Gibco, #26050-070) and penicillin-streptomycin (Gibco, #15140-122) onto 10 cm culture dishes at seeding density of 5-6 x 10^6^ cells/dish and maintained at 37°C in incubation chamber supplied with 5% (v/v) CO_2_ and controlled humidity. Contact media collected from these cultured dishes time to time preserved in -80°C freezers.

Primary mouse hippocampal culture preparation from the E17/18 embryos of time pregnant CD1 dam performed as described in Kaech and Banker, 2006 with modifications (4). The pregnant dam euthanized with Carbon-di-oxide and the pups from uterus extracted in the dissection media. Following the removal of the meninges the bilateral hippocampii carefully isolated from 8-10 pups. The tissue then treated with 2.5% Trypsin for 10 mins at room temperature. Then the tissue washed in the dissection media thoroughly to remove any traces of the residual trypsin. The dissection media then replaced with MEM media supplemented with glucose, penicillin-streptomycin and horse serum. Following this the tissues are made into single cell suspension by passing it multiple times through fire polished Pasteur pipette. The cell concentration then counted using haemocytometer and platted onto 60 mm culture dishes coated with Poly-L-Lysine solution at 5 x 10^5^ cells/dish. This was then maintained in incubation chamber at 37°C supplied with 5% (v/v) CO_2_ and controlled humidity. The media in these dishes were replaced after 8 hours from seeding to Neurobasal (Gibco, #2113049) supplemented with B-27™ (Gibco, #17504001), Glutamine (Sigma-Aldrich, #49419) and penicillin-streptomycin. For long term maintenance, the Neurobasal media further supplemented one tenth of its volume with conditioned media preserved from astroglial feeder culture.

### AAV Transduction of primary neurons *in vitro*

Primary hippocampal neurons cultured in 60 mm dishes were used for experiments involving AAV transduction. The AAV expressing Cerox1 knock-down construct (KD) or universal control (UC) were titre adjusted (1x10^10^ GC/mL) using neurobasal medium. Following this 3/4th of the conditioned media from the days *in vitro* (DIV) 4 culture dishes of primary hippocampal neurons were carefully aspirated into a sterile tube and temporarily stored in refrigerator. The culture dishes then replaced with fresh neurobasal medium with supplements. The diluted AAV preparation then added into the culture dishes and incubated in the tissue culture incubator for 12-16 h. The non-bound virus particles then removed by replacing the contact medium to the initial original medium. The AAV viral transduced culture dishes were then maintained in the tissue culture incubator for 21 days. The total RNA was then prepared from these dishes, using TRIzol™ method for cDNA preparation and quantitative PCR studies. Knockdown constructs expressing AAV virus were transduced (KD1 and KD2) to test the knockdown efficiency in hippocampal neurons. The AAV construct with highest efficiency in the knockdown were used in the stereotaxy based injection into the dorsal hippocampus. AAV preparation, transduction and injections were performed as per approved institutional biosafety (NBRC/IBSC/2022/05 and NBRC/IBSC/2019/06) and animal ethics committee (NBRC/IAEC/2024/193, NBRC/IAEC/2022/179 and NBRC/IAEC/2020/169).

### Stereotaxic injection

Male C57BL/6J animals of 6-8 weeks old were used for the stereotaxic injection delivery of AAV virus into the dorsal hippocampus. The animal was anesthetized with combination of Ketamine (100 mg/kg) and Xylazine (10 mg/kg) administered through intra-peritoneal (IP) injection. The animal’s scalp then carefully shaved to remove the hair and it was mounted on the stereotaxic frame (Neurostar, Drill and microinjection robot) using ear bars and teeth clamp on a heating pad maintained at 37°C. Their scalp was then surface sterilized using liquid povidone solution. An incision was then made on the scalp using surgical blade and extended to reveal the skull beneath. The skull surface was then gently scrubbed with 95% ethanol or 3% (w/v) hydrogen peroxide solution to reveal the Bregma and lambda points. The skull was then carefully drilled guided by surgical microscope (Zeiss, S7 OPMI Sensera®) at the coordinates (Anterio-posterior (AP): -1.9 mm; Medio-lateral (ML): ±1.4 mm) corresponding to the dorsal hippocampus region with reference to the Bregma point. Using a Hamilton syringe (Hamilton, #20750), mounted on to the stereotaxic frame 1 µL of titre adjusted (1-5 x10^8^ TU/ml) AAV vector suspension delivered at Dorso-ventral axis (DV): 1.25 mm depth at the rate of 100 nL/min. Following the injection into bilateral dorsal CA1 region, the scalp skin was proximated and carefully sutured with Monofilament polyamide thread using size 4-0, 3/8 Circle reverse cutting needle (Trusilk, #SN5082). The breathing pattern of the mouse is continuously monitored throughout surgery for any sign of discomfort. The animal was then administered with Meloxicam and Enrofloxacin injections delivered subcutaneously at the dosage recommended by the in-house veterinarian and returned to their original cages. Infrared lamps were used to keep the cages warm for minimum 6 hours following surgery. Post-surgery animals were regularly monitored and applied with topical creams with antibiotic and lidocaine on the suture site for 2-3 days for prevention of infection and pain management.

### Spatial object recognition

Spatial object recognition test performed using male C57Bl/6J mouse of 8 – 12 weeks old. Mice overexpressing Cerox1 or tdTomato were grouped into non-sleep deprived (NSD) and sleep deprived (SD). The animals after 2 weeks of stereotaxic surgery habituated in the arena of spatial object recognition set up for 1 week using gentle cupping method. Following habituation the mice were trained in an acrylic box (80cm x 80cm x 50cm) with simple geometric cues positioned to the walls. Initially the animals were allowed to explore the empty box for 10 mins. The animals were then introduced into the box with the objects (Glass bottle filled with husk and a metallic object positioned diagonally opposite) for learning their position with reference to the geometric cues for 10 mins. Then the animals were returned to their cages and the box was cleaned with 70% (v/v) ethanol. The training was repeated two more times with 5 mins interval to clean the acrylic box. Following the training sessions the animals injected with AAV overexpressing Cerox1 or tdTomato grouped under “SD” were sleep deprived for 5 hours by gentle stroke or disturbing the nest while the “NSD” group animals left undisturbed. After 24 hours the animals tested for their spatial memory by reintroducing into the same box with one of the object displaced from their original position (from diagonally opposite to parallel with reference to the cues). All the activity of the mouse during the training and the testing were recorded using logitech C270 camera with logitech software and saved in .mp4 file format. The amount of time spent by the animals around the objects were tracked and quantified using autotyping 15.4 open source software tool (5). The object preference is then calculated as the ratio of time spent by the animal around the displaced object to the time spent around all objects (6).

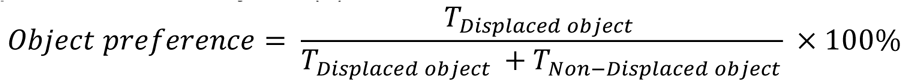

### Mitochondrial Preparation and enzyme assays

The crude mitochondria for the oxidative phosphorylation enzyme assays prepared from the dorsal hippocampus as described previously (7) with modification. For this the animals were euthanized by cervical dislocation, and immediately the dorsal hippocampus region was dissected on ice cold PBS (137 mM NaCl, 2.7 mM KCl, 10 mM Na_2_HPO_4_, 1.8 mM KH_2_PO_4_) solution lacking magnesium and calcium. This was then homogenized in IB1 (225 mM D-Mannitol, 75 mM Sucrose, 0.1 mM EGTA, 20 mM HEPES pH7.4 adjusted with KOH, and 0.5% (v/v) BSA) solution using a pestle homogenizer in an ice bath. The homogenate was then centrifuged at 600 x g for 5 mins at 4°C to remove nuclei and cell debris. The resulting supernatant was carefully aspirated into a fresh tube and centrifuged at 7000 x g for 10 minutes at 4°C. The pellet from this stage was resuspended in ice-cold IB2 (225 mM D-Mannitol, 75 mM sucrose, 20 mM HEPES, 0.5% (v/v) BSA) buffer. This was then centrifuged at 10,000 x g for 10 minutes at 4°C and the resulting mitochondria enriched pellet re-suspended in MRB buffer (250 mM D-Mannitol, 0.5 mM EGTA, 5 mM HEPES). This was then immediately aliquoted and snap-frozen using liquid nitrogen and stored at -80°C deep freezer. The protein content estimated from this fraction using Pierce BCA protein assay kit (Thermoscientific, #23225) as directed by the manufacturer. For the evaluation of Complex I and Complex II enzyme activities, the methodology outlined by (8, 9) was adopted with minor modifications. The crude mitochondrial preparation freeze-thawed before enzyme assays and 50 µg of protein equivalent were utilized to assay the enzyme activity quantification as detailed below in a 96-well titration plate, in a Tecan multimode microplate reader.

The complex I enzyme activity assayed for all crude mitochondria preparations from CA1 region using the oxidation of the NADH (Sigma-Aldrich, # N8129) substrate that exhibit a characteristic change in its absorbance property. For this crude mitochondrial preparation mixed with 25 mM potassium phosphate buffer (pH 7.2), 2.5 mg/mL fatty acid-free albumin (Sigma-Aldrich, # A8806), 5 mM MgCl_2_, 65 µM coenzyme Q (Sigma-Aldrich, #C7956), 2 µg/mL Antimycin A (Sigma-Aldrich, #A8674), and 0.13 mM NADH (Sigma-Aldrich, #N8129). The final volume was adjusted to 0.2 mL using MRB buffer. The reaction mixture maintained at 30°C in the Tecan instrument. The NADH concentration was measured from its absorbance at 340 nm every 5 second intervals for 3 minutes. Following this rotenone (10μM) added into the wells and the absorbance measured for another 2 mins. The rate of NADH to NAD^+^ conversion was determined for both rotenone sensitive and rotenone insensitve period. The specific rate of mitochondrial Complex I activity calculated by subtracting the rate of rotenone sensitive phase to the rotenone insensitive phase. The quantification of Complex II enzyme activity assayed using the DCPIP as a substrate, which exhibits characteristic absorption maxima at 600 nm. The crude mitochondrial preparation was incubated at 30°C for 10 mins in the Tecan instrument mixed with 25 mM potassium phosphate buffer (pH 7.2) and 2 mM sodium succinate. Subsequently, the mixture was combined with 2 µg/mL antimycin, 2 µg/mL rotenone (Sigma-Aldrich, #R8875), and 50 µM Dichlorophenolindophenol (DCPIP) (Sigma-Aldrich, # 33125), with the volume adjusted to 0.2 mL using MRB buffer. The reaction mixture was then measured for absorbance at 600 nm for every 5 seconds over a period of 5 mins, and the rate reduction in the absorbance computed. This reflects the specific activity of the mitochondrial Complex II.

For studies on Cerox1 distribution on mitochondrial matrix pure mitochondria prepared from the crude mitochondria resuspended in the MRB buffer. Crude mitochondrial resuspension from pooled bilateral hippocampii of 4 adult male C57BL/6J animals and carefully layered onto discontinuous Ficoll® gradient (5%-13% w/v in 0.3M sucrose, 10mM EGTA, 10mM Tris) and centrifuged (Beckman coulter, XDR) for 1 hour at 4°C. The free mitochondrial pellet in the bottom was then carefully resuspended in MRB buffer and divided into two fractions. One fraction of the pure mitochondrial suspension subjected to 60μg of RNase A treatment (GCC Biotech, #G4633) at 37°C for 15 mins to remove the mitochondrial membrane associated RNAs while the other fraction remained as a control. Following the treatment RNase traces removed from the mitochondria by washing them thrice in the MRB buffer. The mitochondria then recovered by centrifuging at 20000 x g at 4°C and resuspended in lysis buffer (1% w/v SDS, 10 mM Tris, 10 mM EDTA, 1 mM DTT, 200 U/mL RNase inhibitor). The total RNA isolated from both the fractions using TRIzol reagent and qPCR analysis performed to identify the relative abundance of the Cerox1 distribution normalized using the expression of 18srRNA, 12srRNA and Cox1.

### ATP quantification

Animals were euthanized by cervical dislocation following sleep deprivation and immediately decapitated. The brains were removed and rapidly cooled in ice-cold PBS slurry. The dorsal hippocampus region was then isolated in the ice-cold PBS slurry and homogenized using a pestle in RIPA buffer. The resulting RIPA-solubilized homogenate was aliquoted, and flash-frozen in liquid nitrogen. Samples were then stored at -80°C until further use. The average time from sacrifice to freezing was approximately 3.5 mins.

Protein concentration in the residual homogenate was measured using the BCA assay (Pierce) according to the manufacturer’s instructions. ATP quantification was performed immediately upon thawing the preserved aliquots using a luminescence-based assay kit (Invitrogen, #A22066). For each sample, 10 µg of protein-equivalent RIPA homogenate was mixed in 0.1 mL of RIPA. Assays were carried out in duplicates at two different dilutions against ATP standards prepared in RIPA. Luminescence counts were recorded using a Tecan multimode microplate reader in a white 96-well plate (Thermoscientific™, #136101) with an integration time of 5 s. ATP concentrations were then calculated from standard curve and expressed as relative fold changes, normalized to control ATP levels.

### Statistical analyses

All quantification plots and western blots images are representative of a minimum of three independent experiments. The data are represented in fold change as mean ± SEM. For the analysis of sleep state dependent expression of lncRNA, a standard one-way analysis of variance (ANOVA) used, followed by Bonferroni’s multiple comparisons for post hoc correction. For experiments involving western blot and qPCR analysis, comparing the means of two conditions performed using unpaired two-tailed *t-test* with Welch’s correction. For quantification of the effect of sleep deprivation and Cerox1 overexpression two-way ANOVA performed and the post-hoc analysis performed using Tukey’s method. The analysis of spatial object recognition experiments for the effect of sleep deprivation and the Cerox1 were done using two-way ANOVA method. Bonferroni corrections was performed for comparisons between the means of different groups.

## Notes

### Competing Interest Statement

The authors have declared no competing interest.

